# DeepCAPE: a deep convolutional neural network for the accurate prediction of enhancers

**DOI:** 10.1101/398115

**Authors:** Shengquan Chen, Mingxin Gan, Hairong Lv, Rui Jiang

## Abstract

The establishment of a landscape of enhancers across human cells is crucial to deciphering the mechanism of gene regulation, cell differentiation, and disease development. High-throughput experimental approaches, though having successfully reported enhancers in typical cell lines, are still too costly and time consuming to perform systematic identification of enhancers specific to different cell lines under a variety of disease status. Existing computational methods, though capable of predicting regulatory elements purely relying on DNA sequences, lack the power of cell line-specific screening. Recent studies have suggested that chromatin accessibility of a DNA segment is closely related to its potential function in regulation, and thus may provide useful information in identifying regulatory elements. Motivated by the above understanding, we integrate DNA sequences and chromatin accessibility data to accurately predict enhancers in a cell line-specific manner. We proposed DeepCAPE, a deep convolutional neural network to predict enhancers via the integration of DNA sequences and DNase-seq data. We demonstrate that our model not only consistently outperforms existing methods in the classification of enhancers against background sequences, but also accurately predicts enhancers across different cell lines. We further visualize kernels of the first convolutional layer and show the match of identified sequence signatures and known motifs. We finally demonstrate the potential ability of our model to explain functional implications of putative disease-associated genetic variants and discriminate disease-related enhancers.

## Introduction

Enhancers are distal regulatory elements that can be bound by transcription factors to boost the expression of their target genes. As important regulatory elements, enhancers collaborate with promoters to regulate the transcription of genes in a cis-acting manner, receiving more and more attentions in studies of cell differentiation [1], human diseases [2] and phenotypic diversity [3]. However, due to such facts as far away from target genes, the absence of common sequence features, and the high cell line specificity, it has long been a challenging task to systematically and precisely identify enhancers in a specific cell line.

Enhancers are usually identified by high-throughput biological experiments. For example, Heintzman and Ren [4] used ChIP-seq experiments to establish a landscape of binding sites for individual transcription factors. May et al. [5] mapped binding sites of such transcription coactivators as EP300 and CBP to a large number of enhancers. With the understanding that enhancers are marked by monomethylation of H3K4 [6], genome-wide identification of enhancers have been conducted in large-scale projects such as ENCODE [7] and Roadmap [8]. Besides, using a technique called Cap Analysis of Gene Expression, the FANTOM project [9] has mapped promoters and enhancers that are active in mammalian primary cell lines [10]. Considering that experimental approaches are expensive and time consuming for large scale identification of enhancers, computational methods have been proposed to predict regulatory elements. For example, kmer-SVM used k-mer frequencies of a DNA fragment with a support vector machine to classify regulatory elements [11]. gkmSVM and LS-GKM allowed gaps in a k-mer and improved the prediction performance [12, 13]. Methods based on random forests [14] and decision trees [15] have also been introduced.

Over the past five years, deep learning has been incorporated into bioinformatics studies. For example, DeepBind used a convolutional neural network (CNN) to predict binding proteins and showed higher prediction power than traditional classifiers [16]. DeepSEA learned DNA regulatory codes via a CNN from epigenomic data and predicted effects of noncoding variants [17]. DeepEnhancer predicted enhancers purely relying on DNA sequences and outperformed SVM-based methods [18]. The success of these methods suggests that deep learning is a powerful tool in genomic studies. Nevertheless, all these methods use only DNA sequence information in their models. This formulation, though simple in implementation, obviously lack the power of making predictions in a cell line-specific manner, because DNA sequences are identical in different cell lines.

Chromatin accessibility of the genome has received more and more attentions in the recent years. It is known that putative accessible regions in the genome often work with transcription factors (TFs), RNA polymerases and other cellular machines to regulate gene expression [19]. With the development of high-throughput sequencing techniques, such experimental methods as DNase-seq and ATAC-seq, have enabled the accumulation of a vast amount of chromatin profiles across a variety of cell lines and provides a great opportunity to study transcription factor binding sites (TFBS), DNA methylation sites, histone modification markers, and other regulatory elements [20, 21]. It is therefore natural to integrate DNA sequence and chromatin accessibility information in a single neural network model for the study of cell line-specific enhancers.

With the above understanding, we propose in this paper DeepCAPE, a Deep Convolutional neural network for the Accurate Prediction of Enhancers, using DNA sequences and DNase-seq data. Through comprehensive experiments, we show that our model is not only superior to existing methods in the prediction of enhancers, but also able to predict enhancers across cell lines. With a visualization strategy, we show that sequence motifs discovered by our method successfully match known motifs. Through joint analysis of prediction results with GWAS data, we show the potential ability of our method to identify genetic variants associated with liver cancer and discriminate enhancers related to lymphoma.

## Materials and methods

### Data collection and processing

We used the promoter enhancer slider selector tool (PrESSTo) to download from the FANTOM project experimentally verified enhancers specific to 9 different cell lines, including epithelial cell of esophagus, melanocyte, cardiac fibroblast, keratinocyte, myoblast, stromal cell, mesenchymal cell, natural killer cell and monocyte. We use two strategies to generate negative samples, i.e., non-enhancer fragments that do not overlap with enhancers. First, we randomly sample DNA fragments of variable length from the background genome, with the constraint that the length and GC content of negative samples should be identically distributed as those of known enhancers. The background genome is defined as the entire human reference genome (GRCh37), excluding known enhancers, promoters for coding and noncoding genes, and exonic regions for coding and non-coding genes. Second, we discard the constraint on the GC content to demonstrate the adaptability of our method to different genome contexts. The first model is more stringent and is used throughout this paper. We set the ratios of positive and negative samples to 1:10 and 1:20, i.e., for each positive sample, we generate 10 and 20 negative samples respectively.

We downloaded raw sequencing data of 891 DNase-seq experiments from the ENCODE project [22] and identified experiments corresponding to the cell lines. Given the raw sequencing data of a DNase-seq experiment, we defined the chromatin accessibility score (*S*) of a DNA position as the number of reads (*N*) starting at this position, divided by the average number of reads 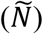 starting at a position in a background region of size *W* centered at the given position. Formally, 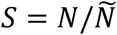 and 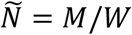 where *M* is the number of reads starting within the background region. A summary of the data is shown in S1 Table.

We consider two issues that are crucial to our method. First, enhancers are of variable length, while a CNN requires inputs of fixed length. Second, a deep neural network has an appetite for a vast amount of training samples. We therefore propose a data augmentation strategy to address both issues (S1 Text).

### Design of DeepCAPE

As illustrated in Fig 1, DeepCAPE consists of four modules. First, a DNA module is used to extract features from DNA sequences. Second, an auto-encoder module is adopted to embed DNase-seq data into a low-dimensional space. Third, a DNase module is used to extract features of chromatin accessibility after dimensionality reduction. Finally, a joint module integrates outputs of the DNA and DNase modules to predict the probability that an input sequence is an enhancer.

**Fig 1.**
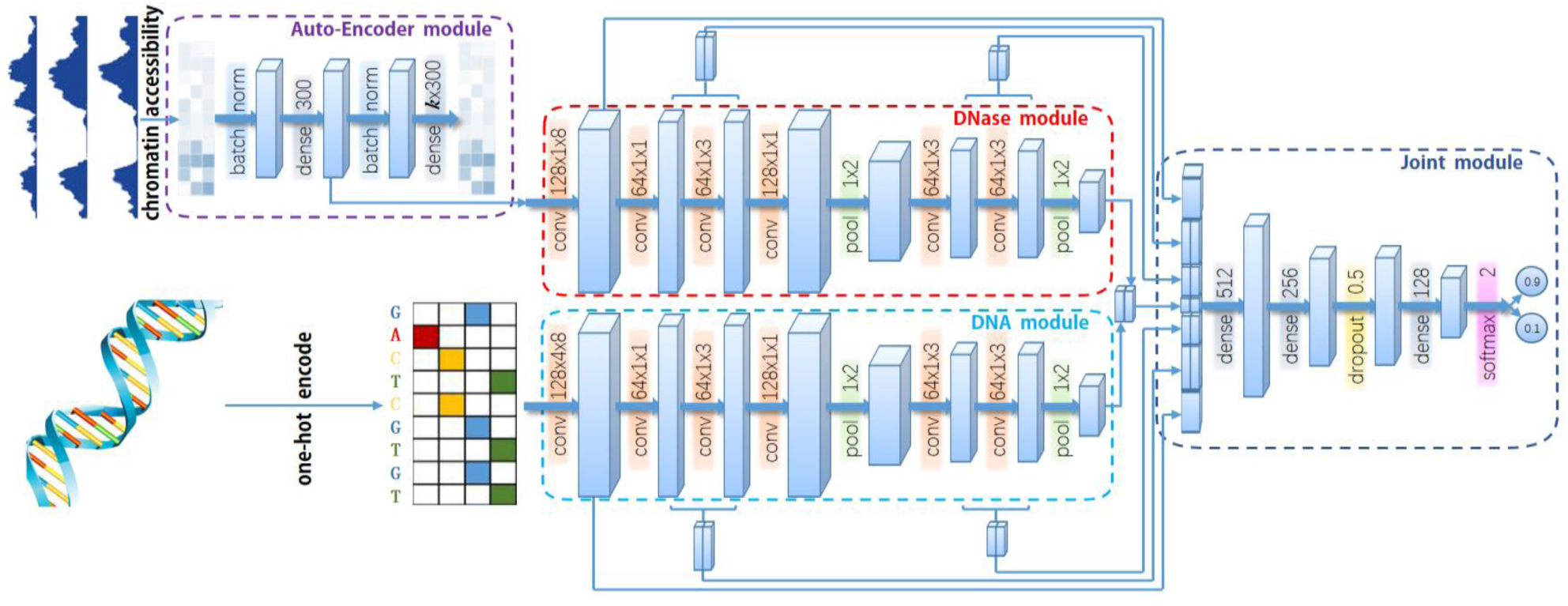
The graphical illustration of DeepCAPE. First, a DNA module is used to extract features from the input DNA fragment. Second, an auto-encoder module is adopted to embed DNase-seq data into a low-dimensional space. Third, a DNase module is used to extract features of chromatin accessibility after dimensionality reduction. Finally, a joint module integrates outputs of the DNA and DNase modules to predict the probability that an input sequence is an enhancer.

#### DNA module

The DNA module is a convolutional neural network (CNN) with multiple convolutional and pooling layers. The first layer uses 128 kernels to scan for sequence motifs of length 8 along the input DNA fragment, which is represented using the one-hot encoding. The second layer uses 64 kernels, each of length 1, to reduce the dimension of features extracted from the first layer by adopting the Network In Network (NIN) model [23], which aims to enhance the discrimination power of the model. The third layer uses 64 kernels, each of length 3, to reduce the number of parameters by drawing on experiences of VGGNet [24]. The fourth layer again adopts the NIN technique and uses 128 kernels, each of length 1, to extract high-level features. The fifth layer adopts the max-pooling strategy to reduce the number of parameters and abstracts features learned in the previous layer. The sixth and seventh layers again adopt the VGGNet technique to further reduce the number of parameters by using 64 kernels, each of length 3. Finally, the eighth layer adopts the max-pooling strategy to abstract final high-level features. In the convolutional layers, the activation of the *k*-th convolutional kernel at the *i*-th position is written as

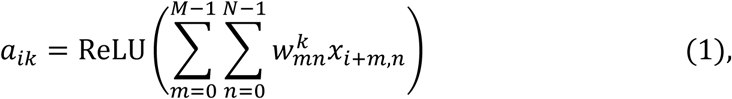

where *X* is the input matrix, *M* the size of the kernel, *N* the number of input channels, 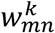 the weight matrix of the kernel. For the first convolution layer, *N* is equal to 4. For other layers, *N* is equal to the number of kernels in the previous layer. The rectified linear unit *ReLU(x) = max(0, x)* activation function sets negative values to zero.

#### Auto-Encoder and DNase modules

A DNase-seq experiment usually has a small number of replicates, and this number varies between cell lines, making the dimensionalities of input data vary between cell lines and preventing the use of a CNN in cross cell line prediction. To solve this problem, we adopt auto-encoder, a neural network designed for unsupervised learning of efficient encodings [25], to embed chromatin accessibility scores of an DNA fragment into a vector of fixed length in a low-dimensional space. Briefly, the auto-encode module first uses a batch-normalization layer to reduce the internal covariate shift and accelerate the training procedure. The output then goes to an encoder component, which is essentially a feedforward neural network that transfers the input data of *k* channels (corresponding to k replicates) into a single channel. After another batch-normalization layer, a decoder component, which is also a feedforward neural network, transfers the data back to k channels. With the module well trained, the decoder is able to produce output similar to the original input, and results of the encoder component can then be used as features extracted from the original data and fed to the successive DNase module. Such an auto-encoder module benefits our model in two aspects. First, regardless of the number of replicates for different cell lines, output of the module is of the same dimension, and thus makes cross cell line prediction possible. Second, effective dimensionality reduction significantly alleviates the computational burden of the successive prediction model.

The DNase module extracts multi-level features from chromatin accessibility scores and is essentially identical to the DNA module in structure, except for the number of input channels. The DNA module is fed with one-hot encoded DNA sequence and has 4 channels, while the DNase module is fed with chromatin accessibility data produced by the encoder component of the auto-encoder module and has a single channel.

#### Joint module

The joint module integrates multi-level features from both the DNA and DNase modules to predict the probability that the input DNA fragment is an enhancer. Drawing on the idea of skip connection in ResNet [26], we merge outputs of the convolutional and max-pooling layers in DNA and DNase modules to form a multi-channel feedforward network. The merged outputs of different layers contain features of different levels, which are integrated via three fully connected hidden dense layers. Such a skip connection strategy endows the model the ability to self-adapt to different sizes of training sets. When there are sufficient training samples, the model may use low-level features. When there are inadequate training samples, the model inclines to explore high-level features automatically. On the top of the architecture, a softmax layer predicts the probability that an input DNA fragment is an enhancer based on the integrated features, as

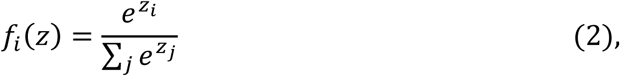

where *f*_*i*_*(z)* is the predicted probability that the input DNA fragment belongs to class *i* (i.e., 1 for enhancer and 0 for non-enhancer).

### Model training

We carry out 5-fold cross-validation experiments to validate the performance of our method for each cell line. Particularly, in order to avoid information disclosure, we partition both positive (known enhancers) and negative (non-enhancers) samples into 5 subsets of nearly equal size before converting sequences of variable length to sequences of fixed length by the data augmentation strategy. In each fold of the experiment, we take 4 subsets to train the model and test its performance using the remaining subset.

Considering that the positive and negative samples are highly imbalanced, we adopt a two-stage training strategy. First, we train an initial model using all positive samples and an equal number of negative samples selected from the training set. After this stage, the DNA and DNase modules obtain the ability to extract features [27]. Then, the joint module is further trained as usual using all the imbalanced samples on the training set, with learning rates of DNA and DNase modules setting to 0. This strategy also alleviates the computational burden. During training, the cross entropy loss is adopted as the objective function to be optimized with Adam (S2 Text).

With a well-trained model, we score all samples augmented from an original sequence on a test set, and then average over these scores to obtain the ?nal probability that the sequence is an enhancer. We also used another strategy that takes the maximum of these scores as the final probability to study the effects of different statistics on the results.

We implement DeepCAPE in Python using Keras [28] with Tensorflow as the backend, while the Theano backend also generated very close results according to our test. The NVIDIA GeForce GTX 1080Ti GPU is used to accelerate the computation.

### Motif visualization

We propose a motif visualization strategy to interpret features extracted by DeepCAPE. We converted kernels of the first convolution layer to probabilistic position weight matrices (PWMs) by counting nucleotide occurrences in the set of sequences that activate the kernels. Briefly, each kernel of the first convolution layer is converted into a PWM by scanning along input sequences for activated positions and then calculating the PWM by pooling corresponding regions [29, 30]. A position *i* is regarded as being activated if

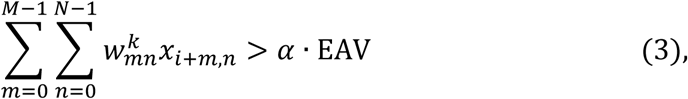

where *α* is the control coefficient (*0 < α < 1*) and EAV the extreme activation value defined as

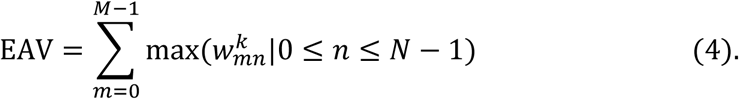

Be set length of kernels in the first convolution layer to 8 and *α* to 0.9. We identify putative sequence motifs by using the tool TomTom 4.11.2 [31] with *q*-value threshold 0.1 to match PWMs identified by our method to the JASPAR database [32].

## Results

### DeepCAPE accurately predicts enhancers

To verify the performance of DeepCAPE, we conducted a series of 5-fold cross-validation experiments using enhancers collected from FANTOM and negative data generated by the background model with the consideration of GC content. We compared the performance of our method with several baseline methods, including gkmSVM, DeepSEA and DeepEnhancer, with parameters proposed by the respective authors. We also proposed a variation of our model, named “seq only”, which discarded the auto-encoder and DNase modules and predicted enhancers using only DNA sequence information. Considering our imbalanced classification task, we computed two widely used metrics, the area under the precision-recall curve (auPRC) and the area under the receiver operating characteristic curve (auROC).

The performance at different ratios of positive and negative samples (1:10 and 1:20) with augmentation stride 1 is shown in Fig 2. Our method consistently outperforms the three baseline methods. In more detail, when the ratios of positive and negative samples are 1:10 and 1:20, respectively, the auPRC scores of our method are on average 0.474 and 0.590 higher than gkmSVM, 0.522 and 0.598 higher than DeepSEA, and 0.511 and 0.588 higher than DeepEnhancer. One-sided paired-sample Wilcoxon rank sum tests consistently suggest that our method achieves higher auPRC scores than a baseline method (*p*-values < 2.2e-16 for all the three baseline methods). In terms of auROC scores, our method is on average 0.121 and 0.151 higher than gkmSVM, 0.169 and 0.151 higher than DeepSEA, and 0.150 and 0.150 higher than DeepEnhancer, when the ratios are 1:10 and 1:20, respectively. Wilcoxon rank sum tests similar to the previous ones also consistently report significant results (*p*-values < 2.2e-16 for all the three baseline methods). All these results suggest the superior performance of our method over existing sequence-based approaches in predicting enhancers.

**Fig 2.**
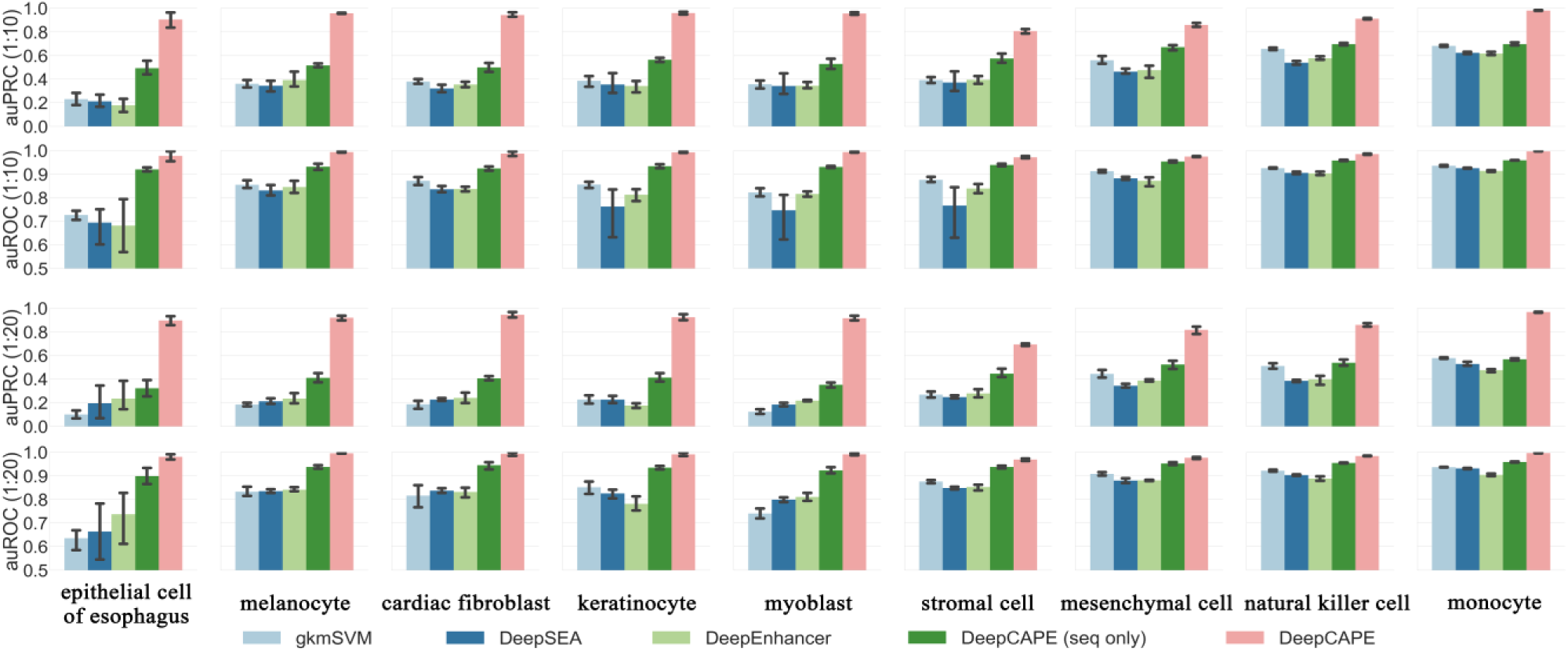
The classification performance measured in auPRC and auROC at different ratios of positive and negative samples (1:10 and 1:20) with augmentation stride 1.

Our method also demonstrates much higher robustness than the baseline methods. With the variance of auPRCs in the 5-fold experiments calculated for each cell line, one-sided Wilcoxon rank sum tests consistently suggest that our method achieves smaller variance than a baseline method (*p*-value=4.019e-4 against gkmSVM, 7.908e-4 against DeepSEA, and 4.571e-3 against DeepEnhancer), suggesting that our method is not sensitive to the partition of training and test samples. Besides, our method consistently performs well in all the cell lines, while the performance of the other three methods shows significant fluctuation across cell lines, suggesting that they are sensitive to the number of training samples.

In terms of model training, benefiting from the usage of dropout layers and the early stop strategy, the performance on the test set is fairly close to that on the training set, indicating that DeepCAPE is able to avoid overfitting. In addition, with regard to the efficiency of model training, DeepCAPE is superior to other deep learning models due to the zero-learning-rate strategy in the second training stage. Take the dataset with augmentation stride 1 of myoblast as an example, when the ratio of positive and negative samples is 1:20, the training time for an epoch is about 126s for DeepCAPE, 301s for DeepSEA, and 237s for DeepEnhancer.

We further conducted a series of experiments to demonstrate the performance of DeepCAPE. First, it is worth noting that the performance of the “seq only” model is also superior to the three baseline methods and performs more steadily, suggesting that our model has the advantage in the case of predicting only with sequences. Second, taking the maximum of the scores of samples augmented from an original test sequence as the final probability generates slightly worse performance, and this may be due to the outliers with high scores in the augmented negative samples. Finally, the performance on datasets without considering GC content is slightly superior to that on datasets under the GC content constraint (S3 Text).

### Contribution of each module

To illustrate the contribution of auto-encoder module, we compared the performance of models using auto-encoder to models without auto-encoder and other two strategies that average the replicates or randomly select a single replicate. As shown in Fig 3 (a), the auto-encoder module not only makes cross cell line prediction possible, but also maintains superior performance of our method even if the dimensionality of the data is reduced (S4 Text).

**Fig 3.**
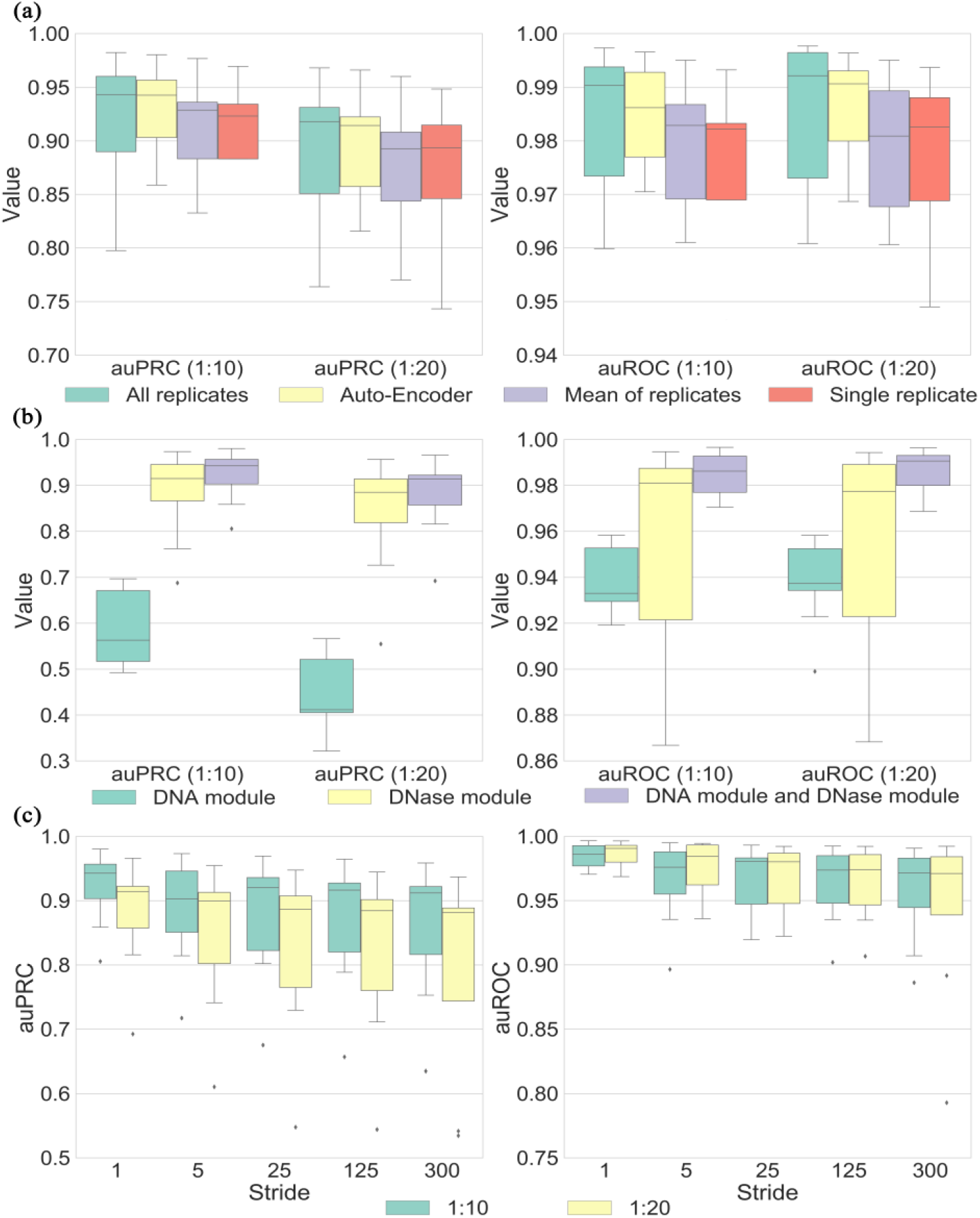
(a) Performance of DeepCAPE with or without the auto-encoder module and other two strategies that average the replicates or randomly select a single replicate. (b) Performance of DeepCAPE with either the DNA or DNase module excluded. (c) Performance of DeepCAPE on datasets of different augmentation strides.

To evaluate contributions of DNA and DNase modules, we performed a model ablation analysis. As shown in Fig 3 (b), DNase-seq data provide more information than DNA sequences to the prediction. In addition, using DNA sequences and DNase-seq data jointly effectively improves performance and stability, indicating that DNA sequences also play an important role in promoting the performance of DeepCAPE and making the performance more stable (S5 Text).

There are more than 100 million parameters in the whole neural network of DeepCAPE, and most of them are concentrated on the merge layer of the joint module. As shown in S2 Fig, we visualized activated features on the merge layer when DeepCAPE was trained with datasets augmented by different strides. With abundant training samples, DeepCAPE is inclined to activate only low-level features, which are extracted by the first three layers. When the sample size is limited, however, DeepCAPE can also activate high-level features, which are extracted by the last three layers (S6 Text).

In order to explore the effect of the number of training samples to the final performance, we repeated the cross-validation experiments on datasets of different augmentation strides for each cell line. As shown in Fig 3 (c), although the performance is gradually decreasing with the augmentation stride becomes longer, the performance is still satisfactory when compared with the three baseline methods and the computational burden is significantly alleviated (S7 Text).

All the observations above suggest that DeepCAPE can not only achieve superior performance with limited known enhancers but also achieve satisfactory performance with longer augmentation strides to effectively save computational time when there are massive enhancers.

### DeepCAPE enables cross cell-line prediction

Experimental approaches are expensive and time consuming for large scale identification of enhancers across a variety of human cell lines. For a cell line whose enhancers have not been identified yet, predicting potential enhancers has great significance in guiding biological experiments for novel enhancers identification.

To accurately predict enhancers across cell lines, we employed a collective scoring strategy. Given a cell line of interest and a DNA fragment, we used models trained on other cell lines to predict the probability that the fragment is an enhancer, and then averaged over these predictions to obtain a final prediction probability. We used the datasets of 9 cell lines from FANTOM to demonstrate the ability of DeepCAPE to predict enhancers in a cross cell line manner. For each cell line, we first excluded the samples that overlap with samples in other cell lines to make sure that there are not common samples with other cell lines. On average, 35.2% and 37.6% of samples are left on the datasets of 9 cell lines when the ratios of positive and negative samples are 1:10 and 1:20, respectively, and the corresponding ratios become 1:8.3 and 1:18.1 averagely. We next used the models of other 8 cell lines to make predictions for the filtered samples of the cell line of interest, and then average over the resulting 8 probabilities to obtain the final prediction probability. We also used other three baseline models to predict enhancers in this cross cell line manner.

As shown in Fig 4, DeepCAPE with our cross cell line predicting strategy is consistently superior to other three baseline methods. In more detail, when the ratio of positive and negative samples is 1:10, the average auPRC and auROC scores of DeepCAPE in 9 cell lines are 0.902 and 0.971, respectively, and when the ratio is 1:20, they are 0.862 and 0.971, respectively. These results suggest that DeepCAPE can accurately predict enhancers across cell lines and thus establish a landscape of potential enhancers specific to a cell line that still lacks systematic exploration of enhancers. The relatively low performance on the dataset of stromal cell may be caused by the fact that we can find only DNase-seq data of stromal cell of bone marrow in ENCODE, which may not match the cell line in FANTOM very well.

**Fig 4.**
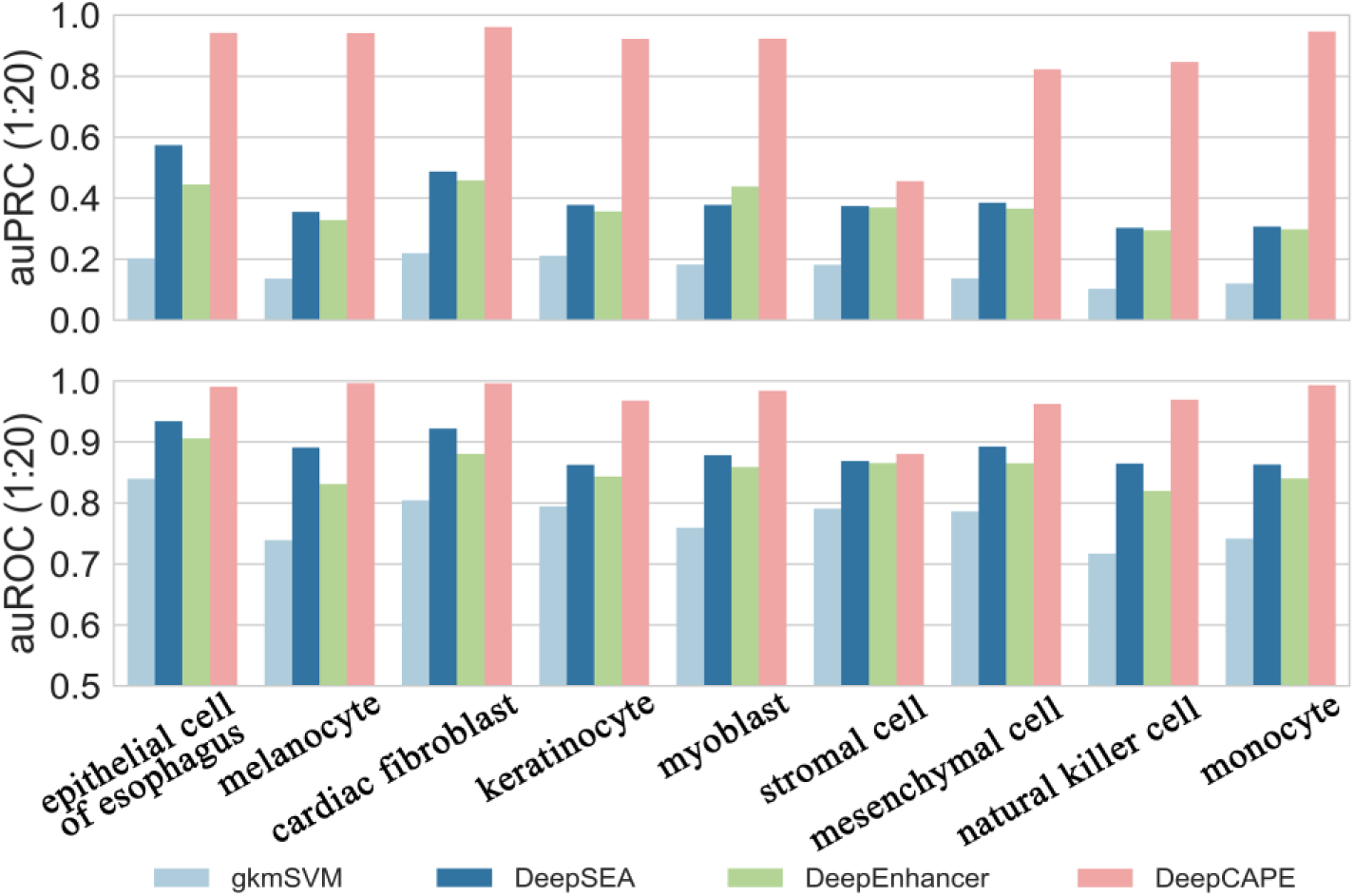
Performance of the cross cell line predicting strategy on 9 datasets from FANTOM. The performance of DeepCAPE is consistently superior to other three baseline methods.

### DeepCAPE recovers known TF binding motifs

To interpret features extracted by DeepCAPE, we used the strategy described in methods to obtain sequence signatures (i.e., PWMs) learned from the first convolution layer of the DNA module. We further identified putative motifs by using the tool TomTom [31] to match these PWMs to the JASPAR database [32].

For each cell line, we displayed the sequence logo of one of the matched motifs in Fig 5. On the dataset of cardiac fibroblast, DeepCAPE recovers SOX21, whose ectopic expression in embryonic stem cells (ESC) induces their differentiation into specific cell types, including those that express markers representative of heart development [33]. In the keratinocyte cell line, DeepCAPE recovers TBX2 which represses transcription from the long control region (LCR) while the composition of the factors binding and regulating the LCR is dependent on differentiation of the host keratinocytes [34]. In the myoblast cell line, DeepCAPE recovers NR4A2 which has been previously shown to contain consensus cAMP response element binding protein (CREB) binding sites that are occupied by CREB and phospho-CREB in myoblasts [35]. On the dataset of natural killer cell, DeepCAPE recovers GATA3 which is a critical regulator for natural killer cell terminal maturation [36]. On the dataset of monocyte, DeepCAPE recovers EGR2, which shows prominent, transient induction in BG-exposed monocytes. It has been demonstrated that enhancers with EGR2 motifs are mainly associated with genes involved in lipid metabolism and biosynthesis and lysosome function [37]. To sum up, DeepCAPE can help us find potential TFs binding in specific cell lines.

**Fig 5.**
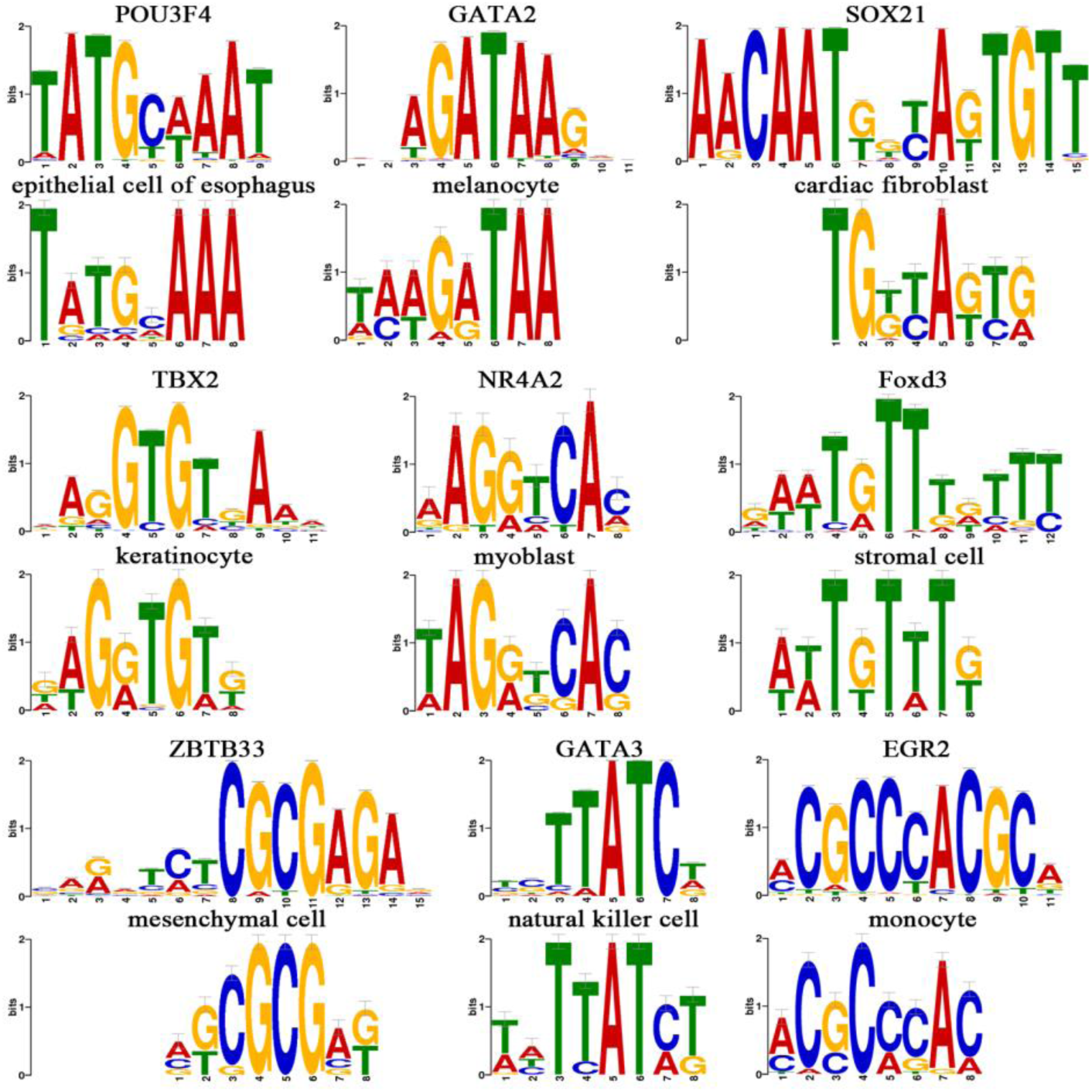
Visualization of motifs learned by DeepCAPE from the first convolutional kernels (above: known motifs from the JASPAR database, below: motifs learned by DeepCAPE).

### Applications of DeepCAPE

To demonstrate potential applications of DeepCAPE, we collected 334 single nucleotide polymorphisms (SNPs) that were possibly associated with liver cancer from GRASP [38]. Each SNP has an association *p*-value obtained from a genome-wide association study (GWAS) regarding liver cancer. We identified a liver cancer cell line (Hepg2) in ENCODE and trained a DeepCAPE model using enhancers and DNase-seq data specific to this cell line. We then calculated a probability that indicated whether a DNA fragment of 300 bps surrounding a SNP was an enhancer for each of the 334 SNPs. We finally classified the SNPs into 5 groups according to logarithmically transformed *p*-values of the SNPs and drew box plots of the predicted probabilities for each group. As shown in Fig 6 (a), predicted probabilities for SNPs with smaller *p*-values are relatively higher than those with larger *p*-values. This observation suggests that predictions given by our method using genomic and epigenomic data are potentially correlated with *p*-values obtained from genetic studies.

**Fig 6.**
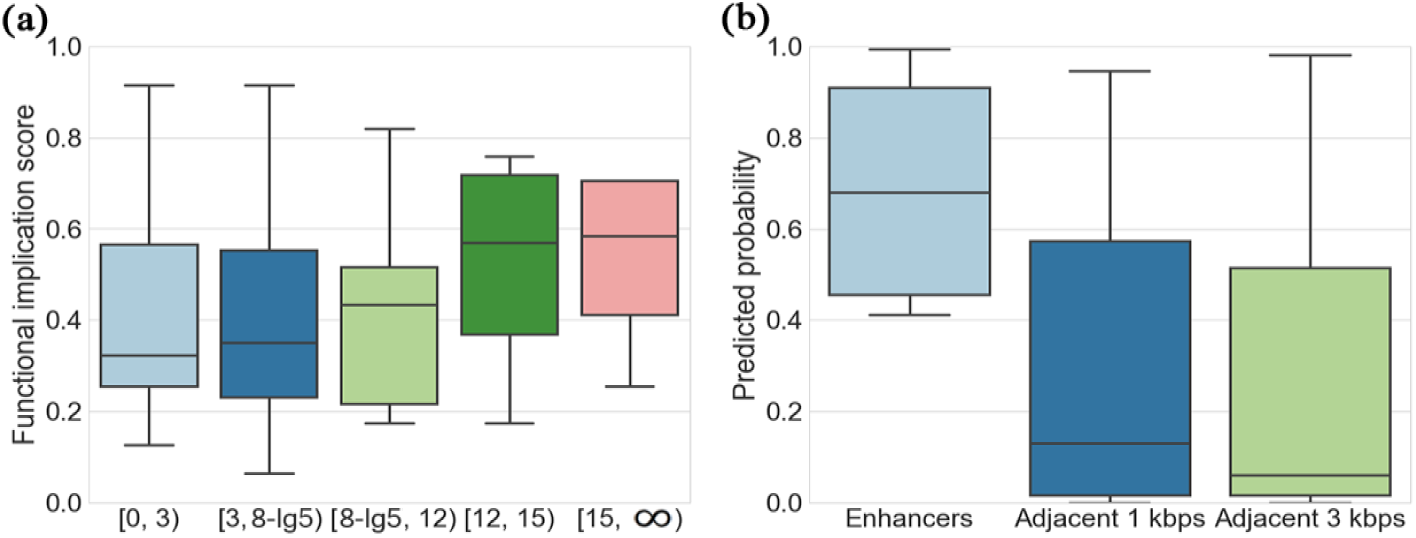
(a) The distributions of predicted functional implication scores of the liver cancer-related SNPs according to different intervals of transformed p-value (-**log**_**10**_***p value***). (b) The distributions of predicted probabilities of the lymphoma-related enhancers and their adjacent sequences sampled from either 1 kbps or 3 kbps upstream and downstream.

We further collected 14 enhancers that are shown to be associated with lymphoma from the literature [39] and showed the ability of our method to discriminant these enhancers from their nearby DNA fragments. For this purpose, we first used enhancers and DNase-seq data specific to a lymphocyte cell line (GM12878) in ENCODE to train a DeepCAPE model. We then used this model to calculate prediction scores for the 14 lymphoma-related enhancers and the same number of their adjacent sequences sampled from either 1 kbps or 3 kbps upstream and downstream. We drew box plot of the prediction scores in Fig 6 (b). It is obvious that prediction scores of the lymphoma-related enhancers are significantly higher than those of the adjacent sequences (one-sided Wilcoxon test *p*-value = 7.915e-6 for adjacent 1 kbps, *p*-value = 9.032e-7 for adjacent 3 kbps). These results suggest that our method has the potential ability to discriminant enhancers linked to lymphoma from their nearby DNA fragments.

## Discussion

We have introduced a deep learning framework named DeepCAPE to integrate DNA sequence information and chromatin accessibility data for predicting enhancers. Through comprehensive validation experiments, we have shown that DeepCAPE is superior to existing methods in a variety of cell lines, capable of making cross cell line predictions, and interpretable in extracted features. We have further demonstrated the potential ability of DeepCAPE to explain functional implications of genetic variants and discriminate disease-related enhancers. Our method has two main application scenarios. First, one can use our method to establish a landscape of potential enhancers specific to a cell line that still lacks systematic exploration of enhancers, thereby promoting the deciphering of regulatory mechanisms for the cell line. Second, one can use our method to explore functional implications of genetic variants or DNA fragments specific to a cell line, thereby bridging genomic and genetic studies towards the understanding of disease development.

Certainly, our work can further be improved in several aspects. First, the incorporation of the long short-term memory (LSTM) network, a kind of recurrent neural network architectures, into our framework may further improve the performance, because LSTM may be able to capture very long-range interaction in the sequence. In addition, the adaptation of an embedding representation of DNA sequences instead of the use of the one-hot encoding may also benefit the prediction accuracy [40]. Second, since we have shown that the first convolutional layer is an effective motif discoverer, researchers may use our model to learn the complex grammar of TF binding in specific cell lines. In addition, one can also explore interactions of motifs in higher convolutional layers. Third, our deep learning framework can possibly be adapted for the identification of other functional elements in the genome, including but not limited to silencers, repressors and insulators. Finally, our framework can also be generalized for the prediction of functional impacts of genomic mutations and the prioritization of candidate variants in whole genome sequencing studies, thereby facilitating both research and practice of precision medicine.

## Acknowledgments

We thank Wanwen Zeng, Wenran Li and Qiao Liu for their helpful discussions. Rui Jiang is a RONG professor at the Institute for Data Science, Tsinghua University.

## Supporting information

### S1 Text

We propose a “fixed-stride” data augmentation strategy as illustrated in S1 Fig. Suppose that the CNN model requires input fragments of length *L*. In the case that a fragment is longer than *L*, we slide a window of size *L* along the original sequence with stride *s* to obtain a number of sequences of length *L*. In the case that an enhancer is shorter than *L*, we slide a window of size *L* along the genome and take sequences overlapping with the original one to obtain augmented sequences. A statistical analysis on a total of 43,011 experimentally veri?ed enhancers in FANTOM shows that the median and mean length of these enhancers are 275 and 288 bps, respectively, and hence we set *L* to 300. We can further control the number of augmented sequences by changing the value of stride *s*. With this strategy, input sequences of variable lengths are converted into ?xed-length samples, and the number of available training samples increase greatly. We also propose an alternative “fixed-ratio” augmentation strategy as follows. Given an original sequence of length *X* and a pre-defined augmentation ratio *r*, we slide a window of size *L* with a derived stride (*L*+*X*)/*r* along the genome to obtain *r* sequences overlapping with the original one. The fixed-stride strategy is used throughout this paper, and the fixed-ratio strategy is used to show the robustness of our model to different data augmentation strategies.

### S2 Text

During training, the cross entropy loss, defined as the entropy between a true distribution *p* and the estimated class probabilities *q*, is adopted as the objective function to be optimized, as

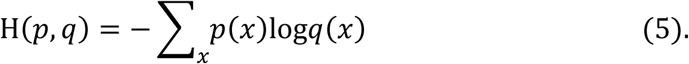

Be adopt Adam, a widely used algorithm for first-order gradient-based optimization of stochastic objective functions [41], to optimize the objective function, with the initial learning rate set to 10-4. The learning rate decay schedule and the early stopping strategy are adopted to accelerate the convergence of training and avoid overfitting. Batch-normalization layers are used to accelerate training by reducing the internal covariate shift [42]. A dropout layer is used between the last two fully connected layers, and it randomly drops half of units to avoid overfitting [43].

### S3 Text

We conducted a series of experiments to further illustrate the performance of DeepCAPE. First, it is worth noting that the performance of the “seq only” model, which only uses DNA sequence information, is also superior to the three baseline methods and performs more steadily, suggesting that our model has the advantage in the case of predicting only with sequences. Second, the “fixed-ratio” data augmentation strategy does not significantly influence the performance of our method. For example, the fixed-stride strategy with stride 1 and the fixed-ratio strategy with ratio 100 produces similar number of augmented sequences, and two-sided paired-sample Wilcoxon tests suggest that their performance is not significantly different (*p*-value = 0.7972, 0.2304, 0.6999 and 0.161 for auPRC (1:10), auROC (1:10), auPRC (1:20) and auROC (1:20), respectively). Third, taking the maximum of the scores of sequences augmented from an original test sample as the final probability generates slightly worse performance with the mean auPRC and auROC decrease by 0.051 and 0.011 respectively when the ratio of positive and negative samples is 1:20. The decline in performance may be due to the outliers with high scores in the augmented negative samples. Finally, we also repeated the prediction experiments with datasets generated by a less rigorous background model without considering GC content to demonstrate the adaptability of our method to different genome contexts. The performance on datasets without considering GC content is slightly superior to that on datasets under the GC content constraint with averagely improved auPRC of 0.041 and auROC of 0.003 when the ratio of positive and negative samples is 1:20.

### S4 Text

Dimensionality reduction of chromatin accessibility scores enabled by the auto-encoder module significantly reduces the amount of data and alleviates the computational burden. To prove that the data after dimensionality reduction is still informative, we repeated for each cell line the 5-fold cross-validation experiments with the auto-encoder module excluded (and thus the dimensions of input data for the DNase-module are different and cross cell line prediction does not work). As shown in Fig 3 (a), the auto-encoder module does not significantly influence the performance (two-sided paired-sample Wilcoxon test *p*-value = 0.5813) but slightly improves the stability of the results. These observations suggest that the auto-encoder module not only makes cross cell line prediction possible, but also maintains superior performance of our method even if the dimensionality of the data is reduced. We also compared the performance of models using auto-encoder to other two strategies that average the replicates or randomly select a single replicate. As shown in Fig 3 (a), the performance of models using auto-encoder is superior to that of averaging the replicates with 2.63% and 3.06% improvement of auPRCs when the ratios of positive and negative samples are 1:10 and 1:20, respectively, and that are 3.59% and 3.56% to the performance of randomly selecting a single replicate.

### S5 Text

We performed a model ablation analysis by repeating the cross-validation experiments with either the DNA or DNase module excluded to evaluate contributions of these modules. As shown in Fig 3 (b), there are evident differences in the contributions of the DNA and DNase modules, especially in terms of the auPRC. After removing the DNA module, the mean auPRCs decrease by 4.64% and 5.84% when the ratios of positive and negative samples are 1:10 and 1:20, respectively. When removing the DNase module, however, the mean auPRCs drop by 36.39% and 49.38%, respectively. Obviously, DNase-seq data provide more information than DNA sequences to accurately predict enhancers. In addition, using DNA sequences and DNase-seq data jointly effectively improves performance and stability, indicating that DNA sequences also play an important role in promoting the performance of DeepCAPE and making the performance more stable.

### S6 Text

As shown in S1 Table and Fig 2, with even only a few thousand training samples, DeepCAPE still performs very well in all the 9 cell lines, while the performance of the three baseline methods is greatly affected by the number of training samples. This means that our method can automatically adapt to different sizes of training sets for better performance and thus achieve superior performance on a dataset with limited number of known enhancers.

We further visualized activated features on the merge layer of the joint module when DeepCAPE was trained with datasets augmented by different strides. With a model trained, we fed positive and negative samples to the network, calculated values of features for each sample, and defined the activation degree of a feature as the absolute difference of its values between positive and negative samples. Taking the cell line of myoblast as an example, we plot heat-maps of activation degrees for features coming from different convolutional and pooling layers in both the DNA and DNase modules in S2 Fig. Briefly, with abundant training samples (e.g., stride 1), DeepCAPE is inclined to activate only low-level features, which are extracted by the first three layers. When the sample size is limited (e.g., stride 300), however, DeepCAPE can also activate high-level features, which are extracted by the last three layers.

### S7 Text

The massive training samples are helpful to improve the prediction performance of our method. However, it may not be necessary to use all the samples for training with the consideration of the computational burden. In order to explore the effect of the number of training samples to the final performance, we repeated the cross-validation experiments on datasets of different augmentation strides for each cell line. As shown in Fig 3 (c), although the performance is gradually decreasing with the augmentation stride becomes longer, the performance is still satisfactory when compared with the three baseline methods. In more detail, we reported in S2 Table the mean auPRC of DeepCAPE in each cell line with different augmentation strides and corresponding time consumed in each epoch when the ratio of positive and negative samples is 1:20. When the stride is 5, the mean auPRC decreases only 4.026%, while 79.753% of the computational time is saved. More extremely, when the stride is 25, the mean auPRC decreases only 6.542% but 96.620% of the computational time is saved.

**S1 Fig.**
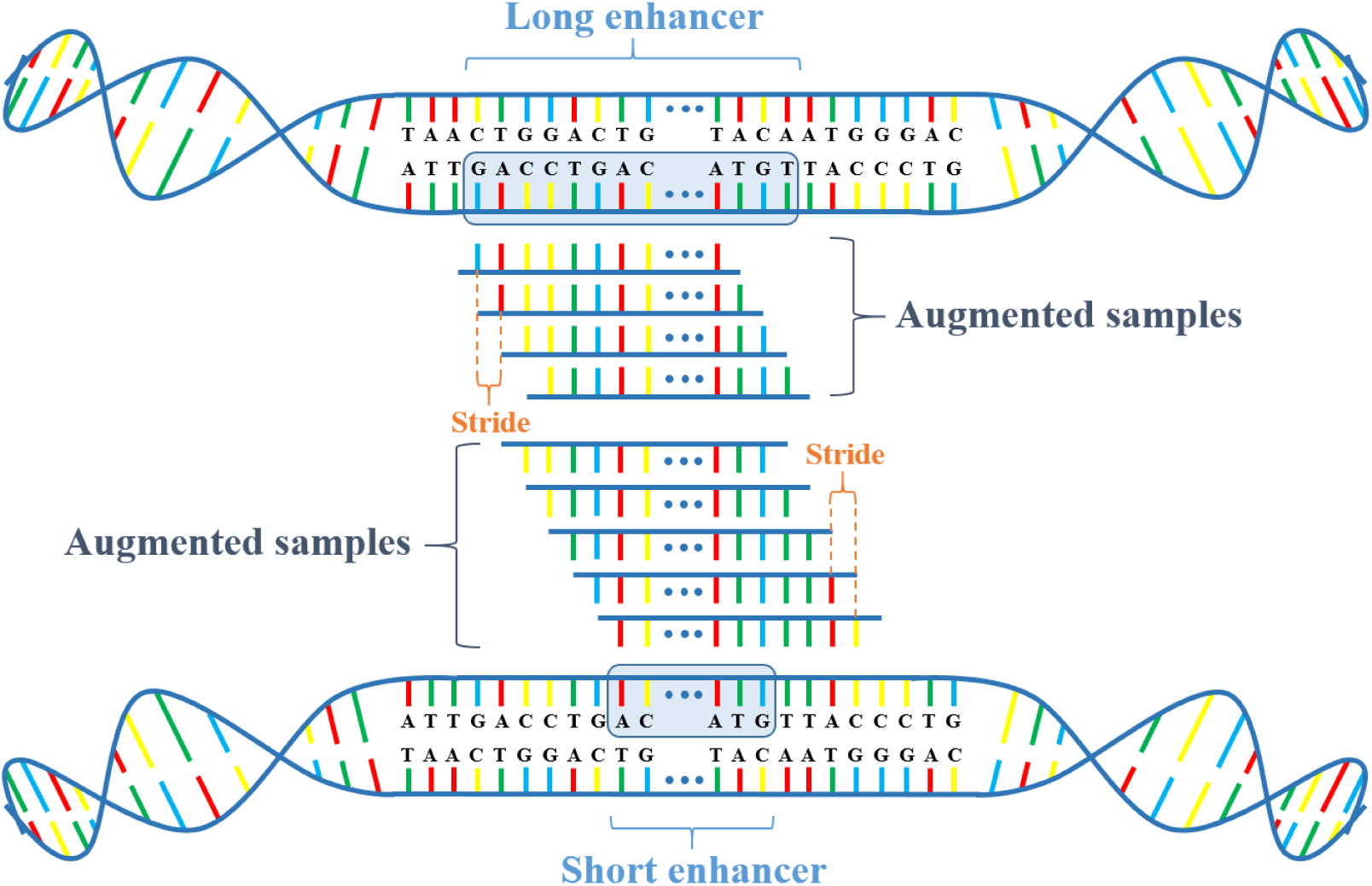
Graphical illustration of the fixed-stride data augmentation strategy. Suppose that input fragments of fixed length *L* is required. When a fragment is longer than *L*, we slide a window of size *L* along the original sequence with stride *s* to obtain a number of sequences with length *L*. When an enhancer is shorter than *L*, we slide a window of size *L* along the genome and take sequences overlapping with the original one.

**S2 Fig.**
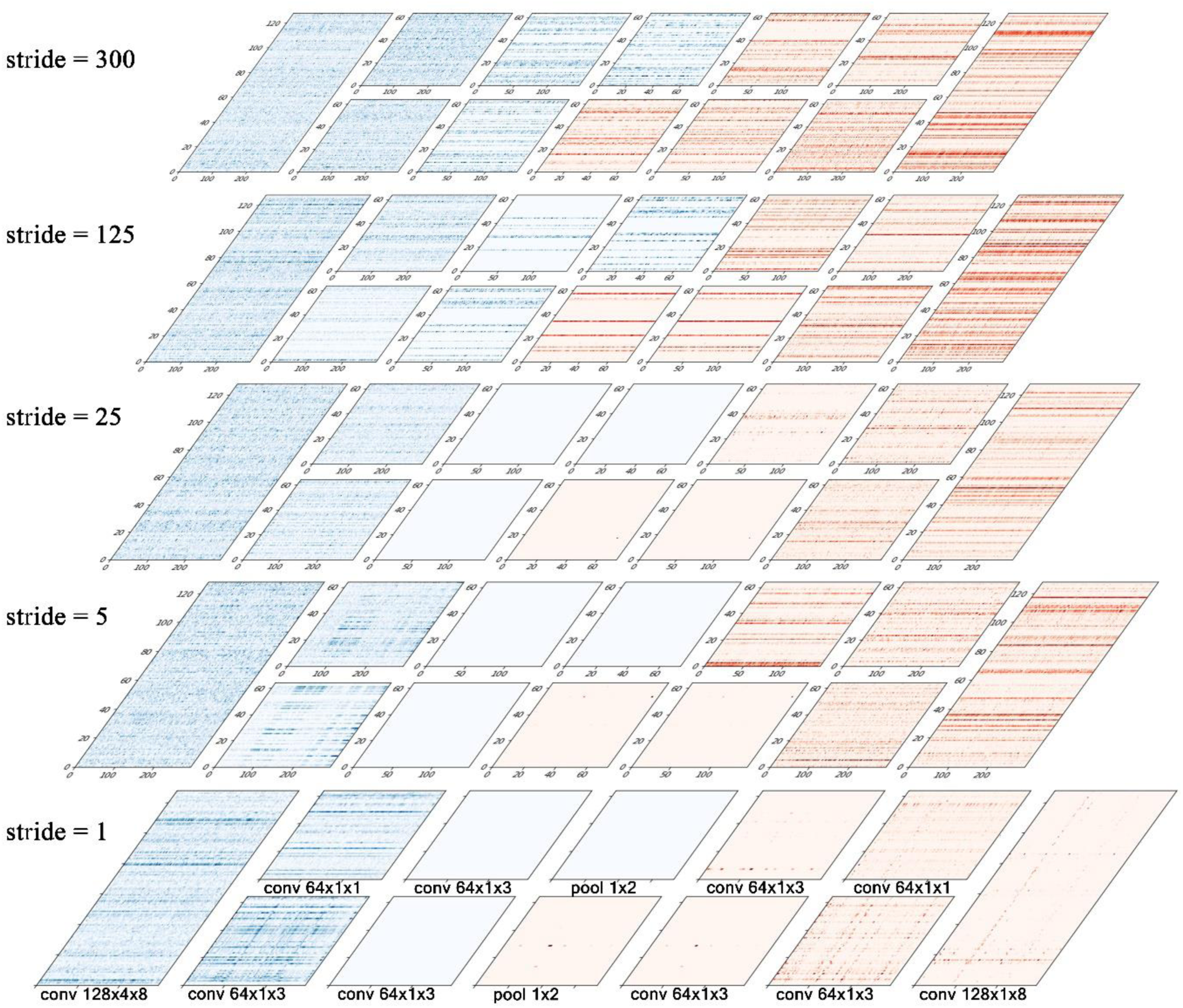
Visualization of activated features on the merge layer extracted by DeepCAPE for training sets of different sizes (different augmentation strides). The blue and red points represent activation degree of features in DNA and DNase modules, respectively (darker color corresponds to higher activation degree). With abundant training samples (e.g., stride 1), DeepCAPE is inclined to activate only low-level features, which are extracted by the first three convolutional layers. When the sample size is limited (e.g., stride 300), however, DeepCAPE can also activate high-level features, which are extracted by the last three layers.

**S1 Table.** Summary of data. Columns are the name of cell line, number of enhancers, number of positive samples after fixed-stride data augmentation (stride 1), and ID of the corresponding DNase-seq experiment.

| Cell Line | Enhancers | Samples | DNase-seq Experiment |
| --- | --- | --- | --- |
| epithelial cell of esophagus | 148 | 15188 | ENCSR000ENN |
| melanocyte | 424 | 45244 | ENCSR518JGY |
| cardiac fibroblast | 446 | 49656 | ENCSR000ENH |
| keratinocyte | 497 | 54343 | ENCSR000EPQ |
| myoblast | 499 | 55238 | ENCSR000EOO |
| stromal cell | 710 | 81295 | ENCSR000EMH |
| mesenchymal cell | 1857 | 215096 | ENCSR405TXU |
| natural killer cell | 2677 | 281512 | ENCSR723JLG |
| monocyte | 7347 | 718064 | ENCSR000EPK |

**S2 Table.** The mean auPRC of DeepCAPE in each cell line with different augmentation strides and corresponding time consumed in each epoch when the ratio of positive and negative samples is 1:20. When the stride is 5, the mean auPRC decreases only 4.026%, while 79.753% of the computational time is saved. More extremely, when the stride is 25, the mean auPRC only decreases 6.542% separately but 96.620% of the computational time is saved, indicating that DeepCAPE can achieve satisfactory performance with longer augmentation strides to effectively save computational time when there are massive enhancer samples.

| Stride | 1 |  | 5 |  | 25 |  |
| --- | --- | --- | --- | --- | --- | --- |
| Cell Line | auPRC | Time (s) | auPRC | Time (s) | auPRC | Time (s) |
| epithelial cell of esophagus | 0.896 | 37 | 0.874 | 7 | 0.857 | 1 |
| melanocyte | 0.919 | 95 | 0.913 | 20 | 0.911 | 3 |
| cardiac fibroblast | 0.943 | 105 | 0.931 | 23 | 0.907 | 4 |
| keratinocyte | 0.922 | 121 | 0.899 | 25 | 0.897 | 4 |
| myoblast | 0.914 | 126 | 0.908 | 25 | 0.887 | 4 |
| stromal cell | 0.692 | 175 | 0.610 | 36 | 0.548 | 7 |
| mesenchymal | 0.816 | 475 | 0.741 | 89 | 0.729 | 17 |
| natural killer cell | 0.857 | 617 | 0.802 | 126 | 0.765 | 22 |
| monocyte | 0.966 | 1631 | 0.954 | 328 | 0.947 | 51 |

